# Ribosomal RNA degradation induced by the bacterial RNA polymerase inhibitor rifampicin

**DOI:** 10.1101/2021.04.24.441238

**Authors:** Lina Hamouche, Leonora Poljak, Agamemnon J. Carpousis

## Abstract

Rifampicin, a broad-spectrum antibiotic, inhibits bacterial RNA polymerase. Here we show that rifampicin treatment of *Escherichia coli* results in a 50% decrease in cell size due to a terminal cell division. This decrease is a consequence of inhibition of transcription as evidenced by an isogenic rifampicin-resistant strain. There is also a 50% decrease in total RNA due mostly to a 90% decrease in 23S and 16S rRNA levels. Control experiments showed this decrease is not an artifact of our RNA purification protocol and therefore due to degradation *in vivo*. Since chromosome replication continues after rifampicin treatment, ribonucleotides from rRNA degradation could be recycled for DNA synthesis. Rifampicin-induced rRNA degradation occurs under different growth conditions and in different strain backgrounds. However, rRNA degradation is never complete thus permitting the re-initiation of growth after removal of rifampicin. The orderly shutdown of growth under conditions where the induction of stress genes is blocked by rifampicin is noteworthy. Inhibition of protein synthesis by chloramphenicol resulted in a partial decrease in 23S and 16S rRNA levels whereas kasugamycin treatment had no effect. Analysis of temperature-sensitive mutant strains implicate RNase E, PNPase and RNase R in rifampicin-induced rRNA degradation. We cannot distinguish between a direct role for RNase E in rRNA degradation versus an indirect role involving a slowdown of mRNA degradation. Since mRNA and rRNA appear to be degraded by the same ribonucleases, competition by rRNA is likely to result in slower mRNA degradation rates in the presence of rifampicin than under normal growth conditions.

## Introduction

RNA is a versatile polynucleotide because of its capacity to form a variety of secondary and tertiary structures (Miao and Westhof 2017). The core translational machinery of life on earth is RNA-based. Ribosomal RNA (rRNA) and transfer RNA (tRNA) mediate decoding and peptidyl transferase activities (Moore and Steitz 2011). Ribosomal RNA and tRNA are highly structured and compact molecules, and interactions with ribosomal proteins, mRNA and enzymes involved in translation contribute to their stability. However, pathways for the degradation of rRNA and tRNA, which eliminate defective molecules and scavenge ribonucleotides during starvation, have been described in *E. coli* (Cheng and Deutscher 2003; Deutscher 2009; Zundel et al. 2009; Svenningsen et al. 2017; Kimura and Waldor 2019; Fessler et al. 2020). Inactive 30S and 50S ribosomal subunits have exposed RNA surfaces, normally buried in the active 70S ribosome, that are targeted for cleavages that initiate the degradation of 23S and 16S rRNA (Basturea et al. 2011; Sulthana et al. 2016). The multienzyme RNA degradosome and the 3’-exo-ribonuclease RNase R, which are involved in these processes, have RNA unwinding activities that facilitate the degradation of structured RNA (Py et al. 1996; Vanzo et al. 1998; Carpousis et al. 1999; Cheng and Deutscher 2005).

Messenger RNA (mRNA) is the molecule that carries protein sequence information. Unlike rRNA and tRNA, the coding regions of *E. coli* mRNA tend to be unstructured and there is a genome-wide correlation between lack of structure and translation efficiency (Burkhardt et al. 2017). Messenger RNA is unstable. In *E. coli*, mRNA half-lives average about 3 minutes, which contrasts with cell doubling times ranging from 20 minutes to greater than 1 hour depending on growth conditions (Bernstein et al. 2004; Esquerre et al. 2014; Chen et al. 2015; Moffitt et al. 2016; Laguerre et al. 2018). Messenger RNA instability is an economical way to ensure that protein synthesis rapidly responds to the reprogramming of transcription in response to changes in growth conditions and stress (Perez-Ortin et al. 2019). Furthermore, mRNA degradation ‘shapes’ the transcriptome since the level of an mRNA coding sequence is a function of both its rate of synthesis and its rate of degradation (Belasco 2017; Chao et al. 2017; Dar and Sorek 2018).

Rifampicin, a potent broad-spectrum antibiotic, is a frontline drug in the treatment of difficult to cure infections such as tuberculosis and leprosy (Alifano et al. 2015). In addition to its medical importance, rifampicin has routinely been used to measure mRNA stability. Rifampicin inhibits transcription of DNA by bacterial RNA polymerase (Campbell et al. 2001). Messenger RNA degradation rates are determined by measuring levels as a function of time after inhibition of transcription by rifampicin (Bernstein et al. 2004; Esquerre et al. 2014; Chen et al. 2015; Moffitt et al. 2016; Laguerre et al. 2018). Since mRNA is unstable, it represents only a small proportion of total RNA, which is composed mostly of rRNA and tRNA (Perez-Ortin et al. 2019). Therefore, it has generally been assumed that total RNA levels are constant after the inhibition of transcription by rifampicin. Here, we report that rifampicin treatment results in a 50% decrease in total RNA level due to the degradation of 23S and 16S rRNA. Rifampicin induced rRNA degradation occurs under different growth conditions and in different strain backgrounds. RNase E, PNPase and RNase R participate in this degradation. The implications of rifampicin-induced rRNA degradation for measurements of mRNA stability are discussed.

## Results

### Cell size decreases after rifampicin treatment

During work on the localization and dynamics of the RNA degradosome of *E. coli* using wide field microscopy, we saw a striking effect of rifampicin on cell size (Fig. 1A). Previous flow cytometry measurements showed that rifampicin treatment reduces cells size (Skarstad et al. 1986). Here, cell dimensions were quantified by direct measurement of 80 to 160 live cells (Fig. 1B and C). The length of the Kti162, SLM018 and SLM024 cells 30 minutes after rifampicin addition was about 60% compared to untreated cells with strong statistical support for this size reduction (P < 0.0001). The effect on cell width is negligible. These strains, which express variants of RNA degradosome components tagged with msfGFP or mCherry, are derivatives of the wild type *E. coli* K12 strain NCM3416.

**Fig. 1.**
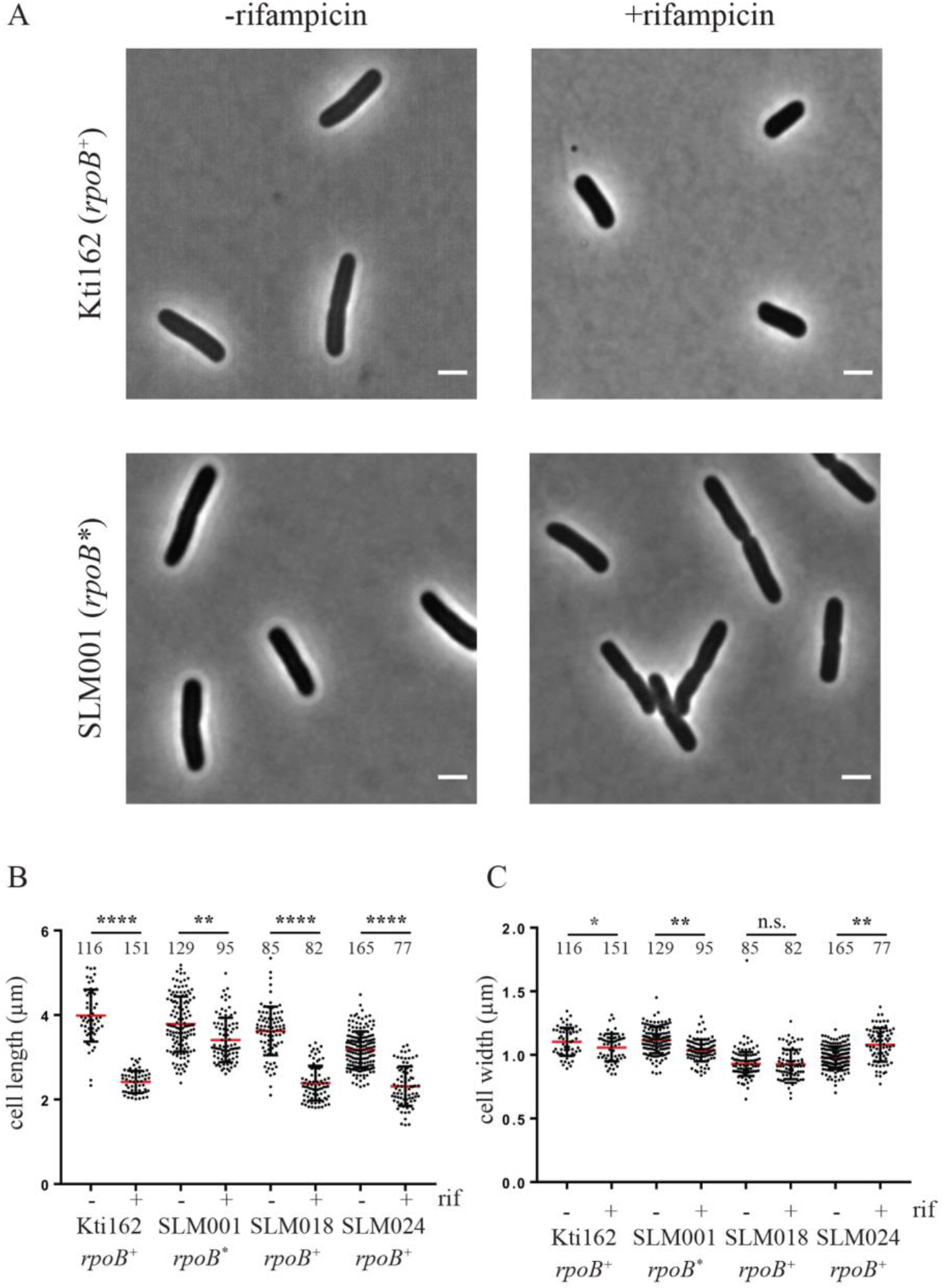
Cell dimensions. The Kti162 strain expressing RNase E-mCherry, SLM018 strain expressing PNPase-msfGFP, SLM024 strain expressing RhlB-msfGFP, and the isogenic rifampicin resistant strain SLM001 were grown to OD_600_ = 0.5 in LB at 37°C. **A**. Phase contrast images before and 30 minutes after adding rifampicin (150 µg/ml). The bar in the lower right corner of each micrograph indicates scale (2 µm). **B. and C**. Scatter plots of cell length (B) and width (C). Since the results of two independent experiments were the same, the data were pooled. The number of cells that were measured are indicated at the top of the distribution profile. Statistical significance of the difference between untreated and rifampicin treated cells was calculated using the non-parametric Mann-Whitney test: **** = P < 0.0001; *** = 0.0001< P <0.001; ** = 0.001< P <0.01; * = 0.01< P <0.05; n.s. = P >0.05.

As a control, we measured cell dimensions in a rifampicin resistant strain (Fig. 1B and C). SLM001, which is isogeneic to Kti162, has an *rpoB(D516Y)* mutation resulting in rifampicin resistant RNA polymerase (Campbell et al. 2001; Alifano et al. 2015). After rifampicin treatment, SLM001 cell length was about 90% of the untreated control (Fig. 1B). Since the SLM001 strain continues to grow in the presence of rifampicin, the small decrease in length is likely due to a slowdown in growth as cells exit exponential growth phase (Akerlund et al. 1995). The SLM001 control shows that the decrease in cell size results from the inhibition of transcription by rifampicin. Since *E. coli* cell shape can be modelled as a cylinder with hemispherical caps, a decrease in cell length of 60% results in a 50% reduction in cell size. This result suggests that cells undergo a terminal reductive cell division after growth is arrested by rifampicin.

### RNA levels decrease after rifampicin treatment

During the isolation of RNA from rifampicin treated cells, we noticed a decrease in yield per ml of culture about 10 minutes after addition of the drug. This was unexpected since mRNA, which is unstable, is only a small proportion of total RNA. Fig. 2A shows the quantification of the levels of total RNA from three biological replicates of rifampicin treated cells. As these cultures were growing exponentially, total RNA levels increase for several minutes after rifampicin addition due to a lag in inhibition of transcription. Remarkably, total RNA levels drop 50% between 5 and 15 minutes after the addition of rifampicin.

**Fig. 2.**
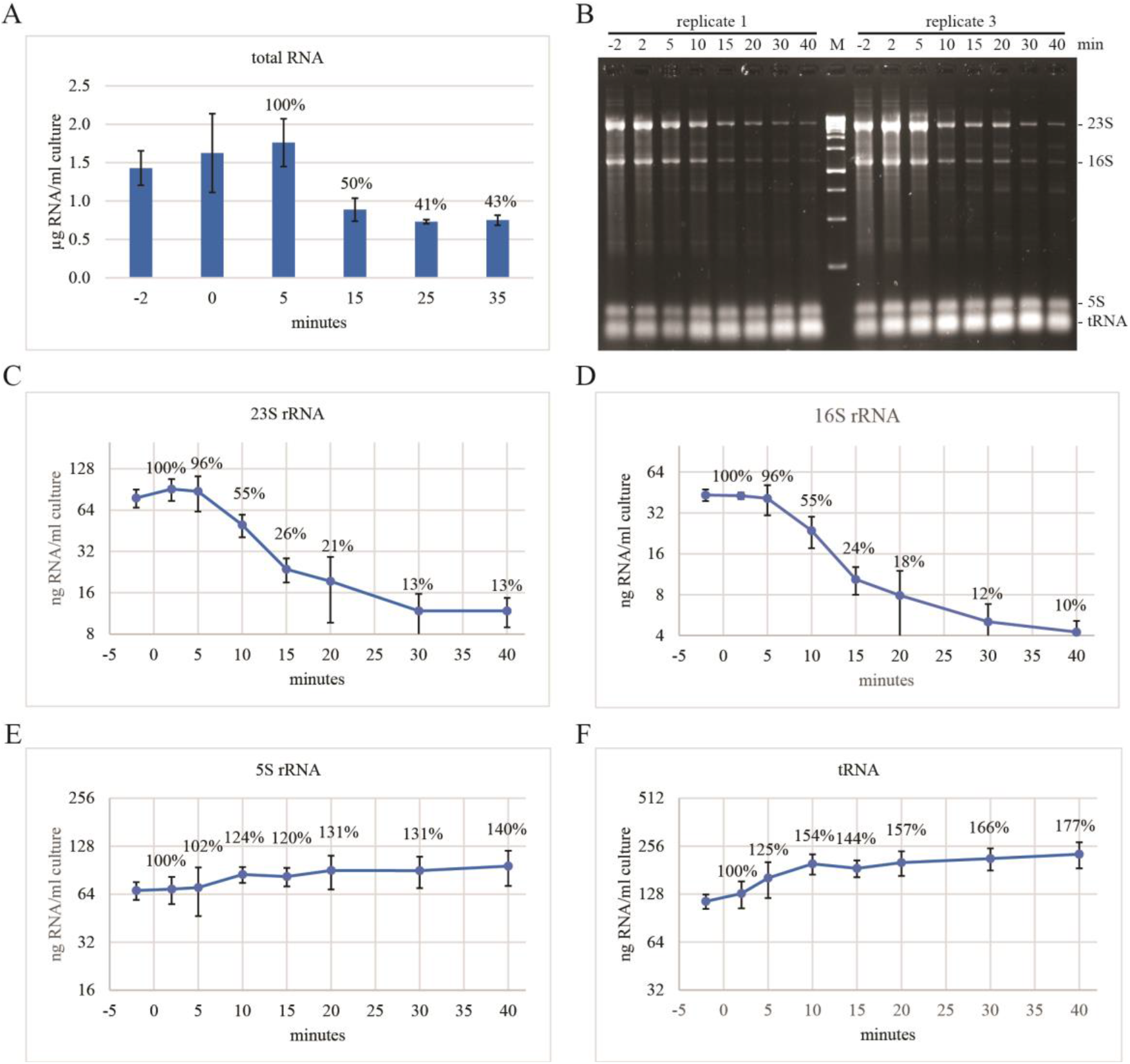
RNA levels after rifampicin treatment. The Kti162 strain was grown in LB at 37°C to OD_600_ = 0.5 and then treated with rifampicin (150 µg/ml). **A**. Total RNA. The 0 min time point was taken immediately after the addition of rifampicin. RNA was extracted from equal volumes of culture. Quantities are expressed as micrograms of RNA per ml of culture. The graph shows the average and standard deviation of three biological replicates. Percentages indicate levels relative to the 5 min. time point. **B**. RNA was fractionated on 2.4% agarose gels and then stained with SYBR™ Safe. A gel loaded with RNA from two biological replicates is shown. M = double-stranded DNA size markers. **C**.**F**. Levels of 23S, 16S and 5S rRNA, and tRNA after rifampicin addition were quantified from three biological replicates. Nominal amounts of RNA were determined using the fluorescence of known quantities of the 500 and 1500 bp double-stranded DNA size markers as standards. The graphs show the average and standard deviation expressed as nanogram of RNA per ml of culture. Percentages indicate levels relative to the 2 min. time point.

To characterize stable RNA species after rifampicin treatment, we separated total RNA by agarose gel electrophoresis, stained the gels with SYBR™ Safe and quantified the levels of 23S, 16S and 5S rRNA, and tRNA (Fig. 2B-F). The quantification shows the average and standard deviation from three biological replicates. There is a sharp 75% decrease in the levels of 23S and 16S rRNA between 5 and 15 minutes after the addition of rifampicin. The levels continue to decrease with time resulting in an approximately 90% decrease by 30 min. The data show that the percent decrease of 23S and 16S rRNA is comparable, which is consistent with degradation of equimolar amounts of each species. In contrast, after an increase in 5S and tRNA levels during the first 10 minutes, which is likely due to the processing of precursors to their mature forms, these RNA species are stable.

The results in Fig. 2B-F show that the loss of 23S and 16S rRNA contributes significantly to the 50% decrease in total RNA after the addition of rifampicin. Note that quantification of levels of total RNA, which is based on UV absorption, cannot be compared directly to nominal levels of individual RNA species, which is based on fluorescence intensities and normalization to known quantities of double-stranded DNA size markers.

Visual inspection of dozens of micrographs such as the ones shown in Fig. 1 gave no indication of cell ghosting or lysis after rifampicin treatment. The conclusion that rifampicin does not disrupt cell integrity is supported by the stability of 5S rRNA and tRNA after rifampicin addition (Fig. 2E-F). Since the RNA extraction protocol employed here involves pelleting cells before lysis in Trizol, this result argues strongly against the possibility that the decrease in 23S and 16S rRNA levels is due to loss of cell integrity and supports the conclusion that cell size reduction is due to terminal reductive cell division.

### Decrease in levels of 23S and 16S rRNA is due to degradation *in vivo*

We performed controls to test the possibility that our results are an artifact of the RNA extraction protocol. RNA was purified using either a Zymo RNA MiniPrep Plus kit or Phase-lock spin tubes and isopropanol precipitation. Fig. 3A shows that the decrease in 23S and 16S rRNA 30 minutes after the addition of rifampicin is the same using either protocol. As a further control, we added purified ribosomes to a Trizol lysate of rifampicin treated cells. Figure 3B, lanes 4 to 6 shows that 23S and 16S rRNA are fully recovered when the Trizol cell lysate (lane 4) was doped with purified ribosomes. Lanes 5 and 6 are controls in which purified ribosomes were extracted with Trizol. These results show that the decrease in 23S and 16S rRNA levels is not due to loss or degradation during the extraction procedure. We therefore conclude that the reduction in levels of 23S and 16S rRNA after the addition of rifampicin is due to degradation *in vivo*.

**Fig. 3.**
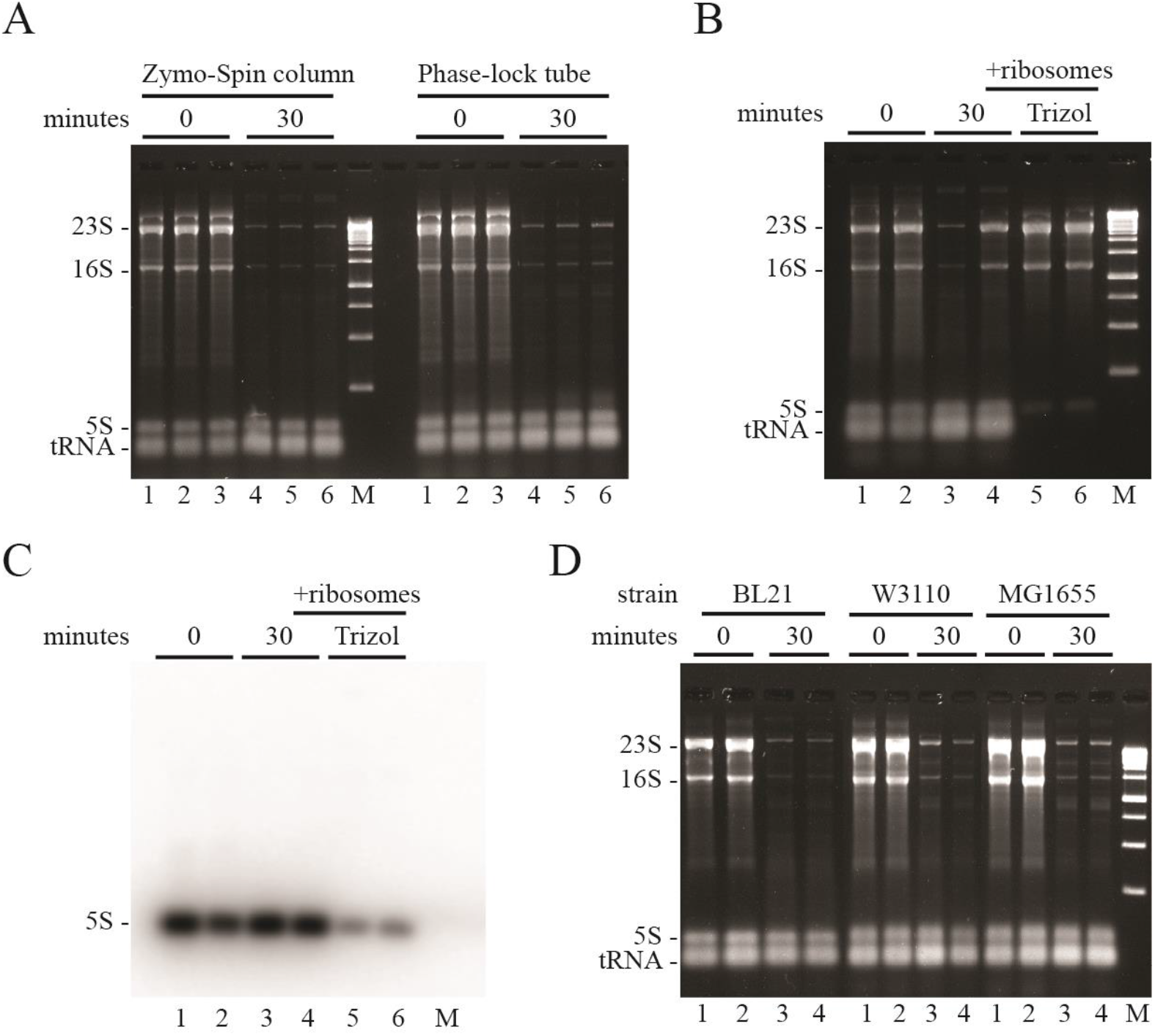
Rifampicin-induced degradation of 23S and 16S rRNA in *E. coli* K12 and B strains. Total RNA extracted from equal volumes of culture was separated by agarose gel electrophoresis. In **A**., **B**. and **D**., the gels were imaged by SybrSafe staining. M = DNA size markers. **A**. RNA was extracted by cell lysis in Trizol and purification with either a Zymo-Spin column kit or Phase-lock fractionation and precipitation with isopropanol. Lanes 1-3, RNA extraction immediately after the addition of rifampicin (biological replicates). Lanes 4-6, corresponding RNA extraction 30 minutes after the addition of rifampicin. **B**. RNA was extracted by cell lysis in Trizol and purification with a Zymo-Spin column kit. Lanes 1 and 2, RNA extraction immediately after the addition of rifampicin (biological replicates). Lanes 3 and 4, corresponding RNA extraction 30 min. after the addition of rifampicin. In lane 4, ribosomes were added to the Trizol cell extract before purification. Lanes 5 and 6, ribosomes were added to Trizol before purification. **C**. Northern blot of RNA from **B**. probed with ^32^P-oligonucleotide specific to 5S rRNA. **D**. Rifampicin-induced rRNA degradation in BL21, W3110 and MG1655 strains of *E. coli*. Lanes 1 and 2, total RNA immediately after the addition of rifampicin (biological replicates). Lanes 3 and 4, corresponding RNA 30 min. after the addition of rifampicin.

The results in Fig. 3B, lanes 1 and 2 vs. lanes 5 and 6 suggest that there is an excess of 5S rRNA *in vivo* relative to 23S and 16S rRNA. The apparent overabundance of 5S rRNA is highly reproducible when RNA is extracted by either of the methods used in Fig. 3. We therefore Northern blotted the RNA in Fig. 3B and probed it with a ^32^P-oligonucleotide specific to 5S rRNA (Fig. 3C). The Northern blot confirms that the species identified by SybrSafe staining is indeed 5S rRNA. The purified ribosomes used in Fig. 3B are a translationally active preparation purchased from New England Biolabs. Visual inspection of the rRNA in Fig. 3B, lanes 5 and 6, suggests that the fluorescence intensities are as expected for a molar ratio of 1 molecule each per ribosome. We therefore conclude that there is more than 1 copy of 5S rRNA per ribosome *in vivo*. This result was not investigated further.

### Rifampicin-induced degradation of rRNA in *E. coli* B and K12 strains

We asked if rifampicin-induced rRNA degradation is characteristic of common lab strains of *E. coli*. BL21 and its derivatives are *E. coli* B strains that are often used as hosts for the overexpression and purification of recombinant proteins. W3110 and MG1655 are *E. coli* K12 strains that have been extensively used in molecular genetics studies. Genome-wide sequence analyses have shown a 92% overlap of coding sequences between the core *E. coli* B and K12 genomes (Studier et al. 2009). There is on average about 1% single nucleotide polymorphisms (SNPs) in coding sequences although the SNPs are not uniformly distributed. NCM3416 is a wild type K12 that has RNase PH activity (*rph*^*+*^). W3110 and MG1655 are *rph*^*-*^ strains and deficient in the metabolism of some sugars and pyrimidine synthesis (Jensen 1993; Soupene et al. 2003). Fig. 3D shows that rRNA degradation is induced by rifampicin in BL21, W3110 and MG1655 at levels comparable to NCM3416 (Fig. 2). This result suggests that rifampicin-induced rRNA degradation is a common characteristic of *E. coli* strains.

### Rifampicin-induced rRNA degradation in MOPS media

We asked if growth media affects rifampicin-induced rRNA degradation. In Fig. 4A-F, MOPS media was supplemented with casamino acids (caa) and three different carbon sources: glucose (glc), succinate (suc) or glycerol (gly). In Fig. 4G-H, growth was with glucose in the absence of casamino acids. Note that the levels of 23S and 16S rRNA depend on growth media, which is consistent with higher ribosome content during faster growth. Under all conditions, there was a significant reduction in 23S and 16S rRNA levels after rifampicin treatment whereas there was no difference in 5S rRNA and tRNA levels. From these results, we conclude that rRNA degradation is induced by rifampicin at fast growth rates in rich medium (LB) as well as at slower growth rates in MOPS synthetic media.

**Fig. 4.**
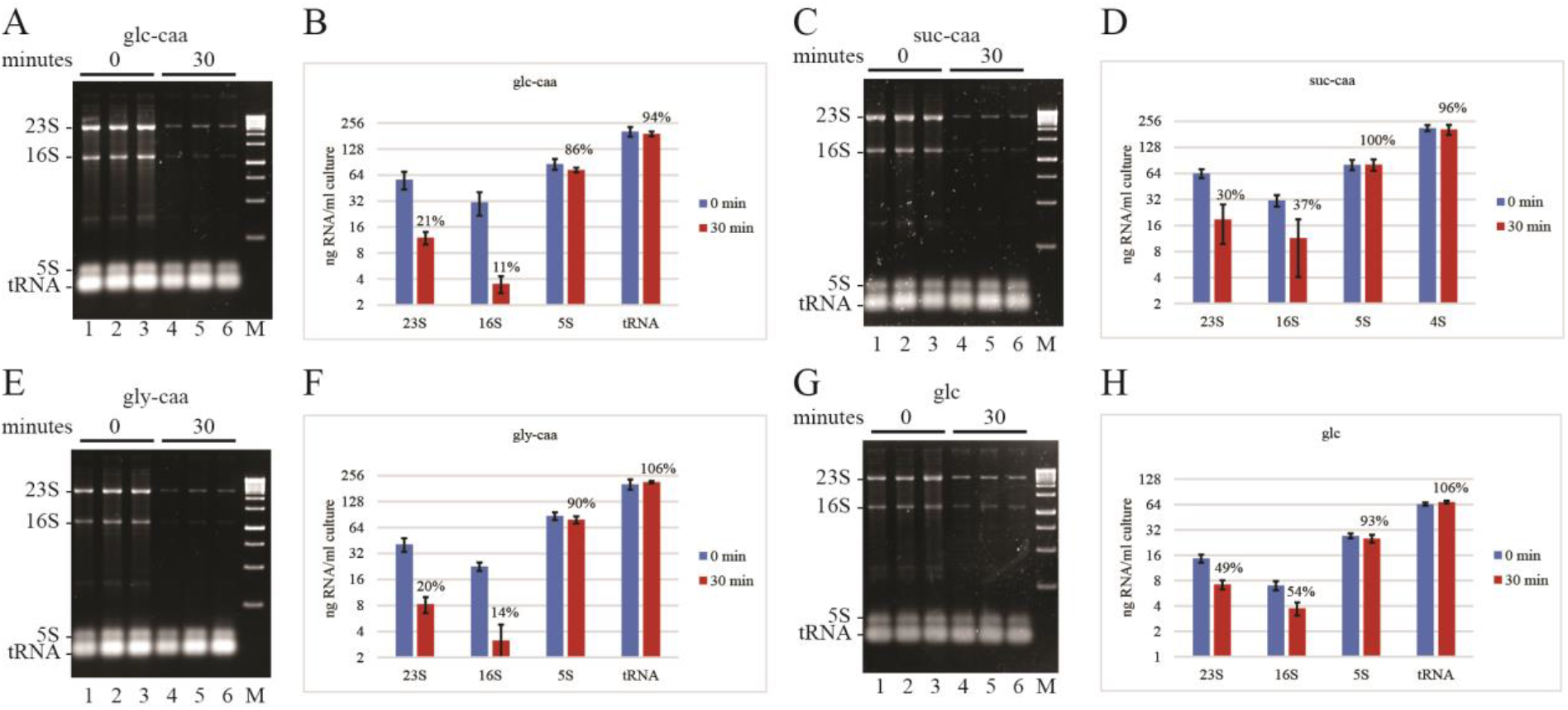
Rifampicin-induced rRNA degradation in MOPS media supplements with carbon sources and casamino acids. **A., C**., **E**. and **G**. SybrSafe stained agarose gels. *E. coli* strain NCM3416 was grown in MOPS medium at 37°C supplemented with glucose (glc), succinate (suc) or glycerol (gly), and casamino acids (caa) except in **G**. Final concentration of carbon source and casamino acids was 0.5% and 0.2%, respectively. Each gel shows three biological replicates. The 0 min time point was taken immediately after the addition of rifampicin. **B., D**., **F**. and **H**. Levels of 23S, 16S and 5S rRNA, and tRNA were quantified from six biological replicates. Nominal amounts of RNA were determined as described (Fig.2). The graphs show the average and standard deviation expressed as nanogram of RNA per ml of culture. Percentages represent normalization of the 30 min. time point after rifampicin addition to the 0 min. time point.

### Cell survival after rifampicin treatment

It is notable that 23S and 16S rRNA are never completely degraded after rifampicin treatment and that final levels of 23S and 16S rRNA are in the range of 5 to 10 ng/ml culture (Figs. 2 and 4). This result suggests that the maintenance of a low level of ribosomes could permit the re-initiation of protein synthesis, and thus cell growth, if the inhibition of transcription is relieved. To determine if rifampicin-treated cells can grow after removal of the antibiotic, we measured colony-forming units (cfu) as a function of time after the addition of rifampicin. Fig. 5 shows a decrease to 46% cfu 5 minutes after rifampicin addition followed by a recovery resulting in a level of 78% after 30 minutes. Since there is a terminal cell division after the addition of rifampicin, the cfu level should increase to 200% by 30 minutes. This result shows that some but not all cells can re-initiate growth after rifampicin treatment. The ability to grow after removal of rifampicin could depend on the number of ribosomes per cell that have escaped degradation (see Discussion).

**Fig. 5.**
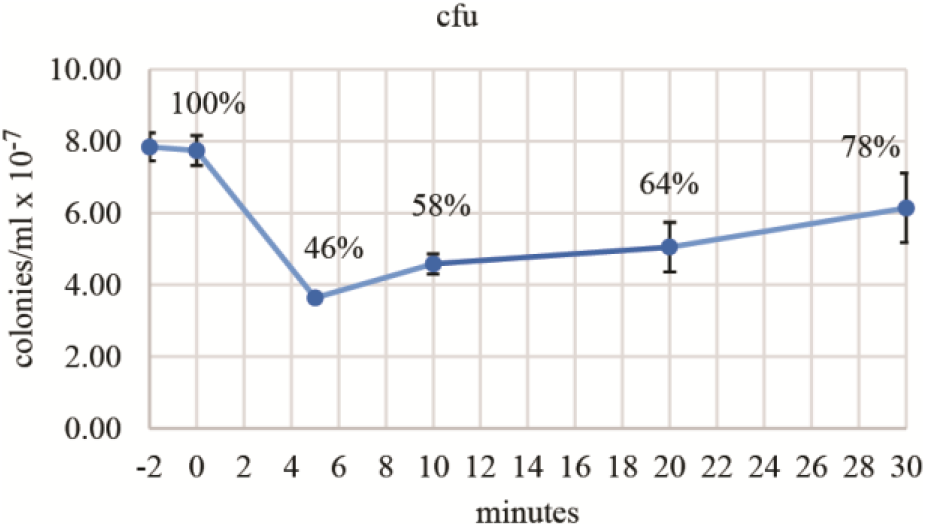
Colony forming units after treatment with rifampicin. Viable cells per ml of culture was measured by colony formation after serial dilution (10^−6^-fold) and plating on LB in the absence of antibiotic. Rifampicin was added at 0 minutes. The graph shows the average and standard deviation of colony forming units (cfu) from three biological replicates. Percentages represent normalization to the -2 min. time point.

### Effect of protein synthesis inhibitors on RNA levels

One consequence of the inhibition of transcription by rifampicin is the inhibition of protein synthesis due to the depletion of mRNA. We therefore tested the effect of chloramphenicol and kasugamycin on stability of 23S and 16S rRNA. Figure 6 shows RNA profiles before and 30 minutes after addition of these inhibitors of protein synthesis. By visual inspection of gel images, we have consistently seen lower levels of 23S and 16S after chloramphenicol treatment. A decrease in 23S rRNA is validated by the quantification in Fig. 6B, but a decrease in 16S rRNA is not statistically significant. We believe that a background smear due to ongoing RNA synthesis interferes with accurate quantification of 16S rRNA levels. Kasugamycin treatment resulted in an increase in 5S rRNA and tRNA levels showing that precursors of these species are processed to their mature form. Stability of the 23S and 16S rRNA suggests that rRNA is not degraded by kasugamycin treatment and that precursors of these species cannot be processed to their mature form. These results show that the inhibition of protein synthesis is not a signal for 23S and 16S rRNA degradation. The result with kasugamycin was unexpected since this antibiotic inhibits translation initiation and we therefore expected that protective interactions between ribosomes and mRNA would be disrupted (see Discussion).

**Fig. 6.**
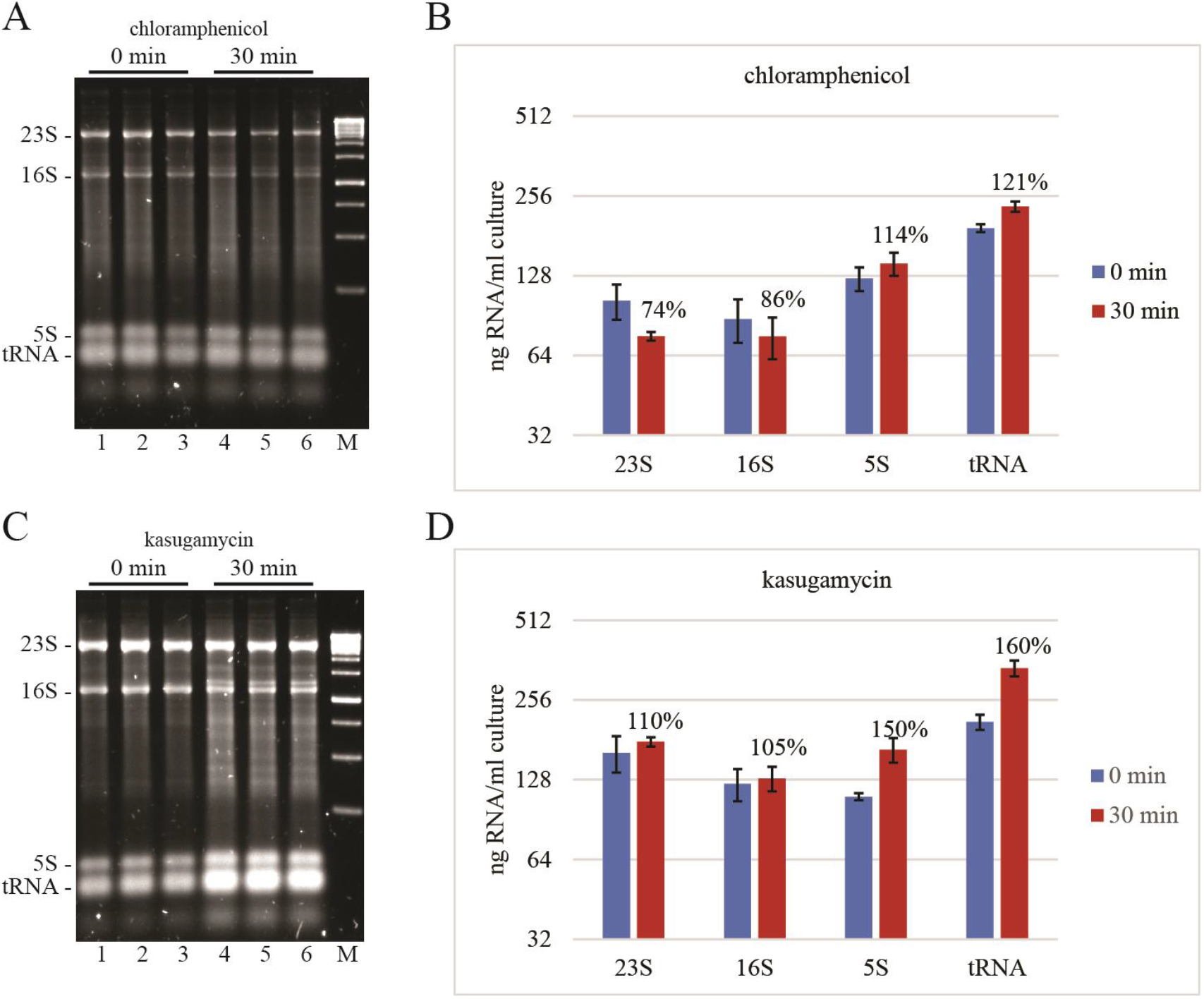
Effect of protein synthesis inhibitors on rRNA and tRNA levels. Total RNA extracted from equal volumes of culture was separated by agarose gel electrophoresis. The gels were imaged by SybrSafe staining. M = DNA size markers. **A. and C**. NCM3416 grown in LB medium at 37°C was treated with chloramphenicol (125 µg/ml) or kasugamycin (1 mg/ml). The 0 min. time points were taken immediately after addition of antibiotic. Lanes 1-3 are biologicals replicates; lanes 4-6, the corresponding 30 min. time points. **B. and D**. Quantification of 23S, 16S, 5S rRNA, and tRNA levels showing the average and standard deviation from six biological replicates. Percentages represent normalization to the 0 min. time point.

### Ribonucleases involved in rifampicin-induced rRNA degradation

To identify the enzymes involved in the degradation of 23 and 16S rRNA, we analyzed strains with mutations that inactivate genes involved in RNA processing and degradation. Fig. 7A and B show the result of rifampicin treatment of nine strains that are lacking activity of the following enzymes: RNase I, RNase II, RNase G, RNase R, RNase PH, poly(A)polymerase, PNPase, YbeY and RNase III. Visual inspection of the gels shows that rRNA degradation is induced by rifampicin in each of these strains thus suggesting that they are not involved in rifampicin-induced rRNA degradation.

**Fig. 7.**
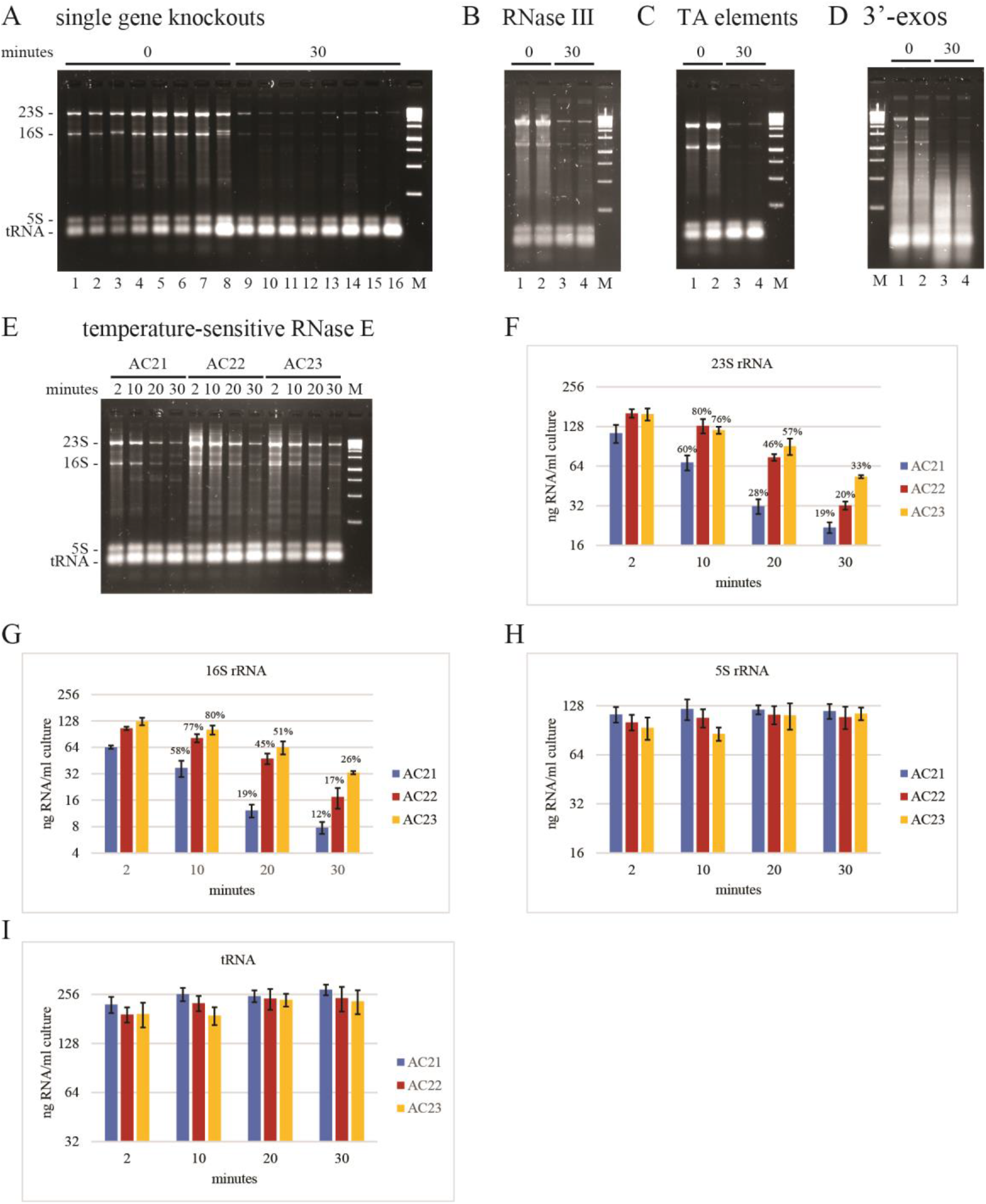
Ribonucleases involved in rifampicin-induced rRNA degradation. Total RNA extracted from equal volumes of culture was separated by agarose gel electrophoresis. The gels were imaged by SybrSafe staining. M = DNA size markers. In panels **A**. to **D**., the 0 min. time points were taken immediately after the addition of rifampicin (150 µg/ml). **A. Single gene knockouts**. Lanes 1 to 8 correspond to mutant strains with gene deletions that result in the loss of the following proteins: RNase I, RNase II, RNase G, RNase R, RNase PH, poly(A)polymerase, PNPase and YbeY. **B. RNase III**. Rifampicin treatment of the BL321 strain, which contains the *rnc105* mutant allele resulting in a lack of RNase III activity (Portier et al. 1987). **C. TA elements**. Rifampicin treatment of the ΔTA10 strain, in which ten toxin-antitoxin elements have been disrupted (Goormaghtigh et al. 2018). Each of these elements encodes an mRNA interferase that inactivates mRNA by endoribonucleolytic cleavage. **D. 3’-exos**. Rifampicin treatment of the CA244 strain, which contains a knockout of the *rnr* gene encoding RNase R and the mutant *pnp200* allele encoding a temperature-sensitive variant of PNPase (Cheng and Deutscher 2003). The strain was grown to OD_600_ = 0.1 at 30°C in LB and then shifted to 42°C. Rifampicin was added 4 hours after the temperature shift. **E. RNase E**. AC21, AC22 and AC23 are isogenic strains containing the *rne*^*+*^, *rne(F68L)*^*ts*^ and *rne(G66S)*^*ts*^ alleles, respectively (Carpousis et al. 1994). The strains were grown to OD_600_ = 0.5 at 30°C in LB and then shifted to 43.5°C for 10 minutes before the addition of rifampicin (150 µg/ml). RNA was extracted 2, 10, 20 and 30 minutes after the addition of rifampicin. **F. - I. Quantification of RNA levels**. The bar graphs show the average and standard deviation of 23S, 16S, and 5S rRNA, and tRNA levels from three biological replicates performed as shown in panel **E**. AC21 is the isogenic wild type control. AC22 and AC23 harbor mutant alleles encoding temperature-sensitive variants of RNase E. Percentages represent normalization to the 2 min. time point.

mRNA interferases are encoded by many toxin-antitoxin (TA) elements in the *E. coli* genome. These enzymes inactivate mRNA by endoribonucleolytic cleavage. In principle, rifampicin treatment should induce mRNA interferase activity since the synthesis of anti-toxins, which are short-lived, is inhibited. We therefore tested the possibility that these endoribonucleases are involved in 23S and 16S rRNA degradation. The Δ10TA strain was engineered to delete ten TA elements encoding mRNA interferases including MazF, RelB, HicA, HigB and MqsR (Goormaghtigh et al. 2018). The gel in Fig. 7C shows that rifampicin induces rRNA degradation in the Δ10TA strain thus suggesting that mRNA interferases encoded by these TA elements are not involved in rRNA degradation.

RNase R and PNPase have been implicated in rRNA quality control and rRNA degradation during starvation for a carbon source (Cheng and Deutscher 2003; Deutscher 2009; Zundel et al. 2009). These enzymes, which are major 3’-exoribonucleases in RNA turnover, have overlapping substrate specificities. Disruption of both genes encoding these enzymes is lethal. We therefore tested a temperature-sensitive strain that is defective for RNase R and PNPase activity at 42°C. In Fig. 7D, as expected, there is a large smear of low molecular weight RNA fragments in the 0 minute time point due to the lack of 3’-exoribonuclease activity. Thirty minutes after the addition of rifampicin, the levels of 23S and 16S rRNA decreased and the level of low molecular weight fragments increased. This result is consistent with a pathway for rifampicin-induced rRNA degradation in which rRNA fragments are degraded by RNase R and PNPase.

We next turned our attention to RNase E, an essential endoribonuclease in *E. coli*, that has been implicated in rRNA degradation upon starvation for carbon (Basturea et al. 2011; Sulthana et al. 2016). We tested two temperature-sensitive strains (AC22 and AC23) in which RNase E is inactivated upon a shift from 30°C to 43.5°C. AC21 is the wild type control. Since mRNA degradation at the non-permissive temperature is slowed but not abolished in these strains (Ono and Kuwano 1979; Babitzke and Kushner 1991), we performed the time courses shown in Fig. 7E. The quantification in Fig. 7F-I shows that there is no difference in 5S rRNA and tRNA levels after rifampicin addition. The apparent higher levels of 23S and 16S rRNA in the temperature-sensitive strains is due a background of undegraded RNA that accumulates after the shift to 43.5°C. There is clear lag in the degradation of 23S and 16S rRNA in the temperature-sensitive strains that is coherent with visual inspection of the gel in Fig. 7E. We therefore conclude that RNase E cleavage is involved in rifampicin-induced degradation of rRNA.

The results in Fig. 7 show that rifampicin-induced rRNA degradation involves the activities of RNase E, PNPase and RNase R. The same pathway has been described for rRNA quality control and rRNA degradation upon carbon source starvation (Basturea et al. 2011; Sulthana et al. 2016). RNase E is believed to initiate rRNA degradation by nicking inactive ribosomal subunits. However, our results raise the issue of whether RNase E acts directly in rRNA degradation or indirectly since a slowdown of mRNA degradation in the temperature-sensitive strains results in a lag in the accumulation of inactive ribosomal subunits (see Discussion).

## Discussion

The arrest of growth by rifampicin is a potentially catastrophic event since inhibition of transcription prevents stress responses that require the synthesis of new RNA. For example, the entry of *E. coli* into stationary phase involves RpoS-dependent transcription and many stress responses require the induction of sRNA synthesis (Battesti et al. 2011; Hor et al. 2020). By imaging live cells on agarose pads and measuring their dimensions, we have shown a 50% reduction in cell size within 30 minutes of the addition of rifampicin, which is likely due to a terminal cell division. The initiation of chromosome replication is a tightly regulated and synchronized process that occurs once per cell division cycle (Skarstad and Katayama 2013). Drugs that inhibit transcription (rifampicin) and protein synthesis (chloramphenicol) block the initiation of chromosome replication. However, DNA synthesis is not inhibited and ongoing chromosome replication continues to completion. Since rapidly growing cells during exponential growth in rich medium divide faster than the time it takes to replicate a chromosome, there are more than two replication forks per cell. Due to completion of DNA replication, measurements showed that over 80% of cells had four chromosomes 90 minutes after the addition of rifampicin (Skarstad et al. 1986). The reduction in cell size together with the increase in chromosome copy number suggests significant remodeling of the cell with a large reduction in the ratio of cytoplasm to nucleoid (Gray et al. 2019). Indeed, single cell measurements have shown that the nucleoid fully occupies the interior of the cell after rifampicin treatment (Bakshi et al. 2014). Taken together, these results show that rifampicin treatment resets the cell cycle to a resting state that preserves chromosome integrity. It is noteworthy that this reset occurs under conditions where transcription is blocked by rifampicin.

We have shown that there is a striking 50% decrease in total RNA after rifampicin treatment of cultures of *E. coli* in exponential growth phase. Electrophoretic separation of total RNA revealed an 90% decrease in 23S and 16S rRNA levels whereas 5S rRNA and tRNA levels were stable. We performed controls showing that the decrease in 23S and 16S rRNA is not an artifact of our RNA extraction protocol and therefore conclude that the decrease is due to degradation *in vivo*. We have observed rifampicin-dependent rRNA degradation in *E. coli* B and K12 strains as well as under different growth conditions. From these results we conclude that rifampicin-induced rRNA degradation is a general property of *E. coli* that occurs over a wide range of growth conditions and that does not depend on carbon source or amino acids. Our results suggest that nucleotides from rRNA degradation are recycled to deoxyribonucleotides for DNA synthesis. We estimate that the degradation of 2000 copies of 23S and 16S rRNA is sufficient for the synthesis of 1 chromosome.

An overview of the data in Figs. 2 and 4 shows that 23S and 16S rRNA degradation is never complete. Final levels are in the range of 5 to 10 ng/ml of culture, which is 10-fold lower than the levels in cells growing exponentially in rich medium. The results of cfu measurements in Fig. 5, which show that approximately 40% of cells 30 minutes after the addition of rifampicin can form colonies, suggest that a basal level of functional ribosomes contributes to the renewal of cell growth if the block in transcription is removed. The number of ribosomes per cell after rifampicin treatment could vary if rRNA degradation is a stochastic process. The capacity to form a colony could therefore depend on a threshold above which there are enough ribosomes to support the level of protein synthesis needed to re-initiate cell growth.

Chloramphenicol treatment resulted in a small reduction of 23S and 16S rRNA levels. Since chloramphenicol inhibits translation elongation (Lambert 2012), this result suggests that ribosomes blocked on mRNA are at least partially protected from rRNA degradation. The continued synthesis of mRNA after chloramphenicol treatment could contribute to this protection. Kasugamycin treatment resulted in an increase in 5S rRNA and tRNA levels showing that precursors of these species can be processed to their mature forms in the presence of this antibiotic. There was no change in 23S and 16S rRNA levels. The apparent stability of 23S and 16S rRNA was unexpected since kasugamycin inhibits the initiation of translation of mRNA (Muller et al. 2016). We expected that the accumulation of inactive ribosomes due to the block in translation initiation would result in rRNA degradation. There is a low level of protein synthesis after addition of kasugamycin that has been attributed to the inability of this antibiotic to block translation of leaderless mRNA (Muller et al. 2016). Therefore, ribosomes could form a protective interaction with mRNA (or mRNA fragments) in the presence of kasugamycin. An alternate but seemingly unlikely possibility is that mature 23S and 16S rRNA is degraded and then replaced by the processing of newly synthesized precursors to their mature form. Regardless of these considerations, the inhibition of translation and cell growth by chloramphenicol and kasugamycin does not provoke the massive decrease in 23S and 16S rRNA levels after rifampicin treatment.

Ribosomal RNA is degraded under condition such as glucose starvation. Considering experimental evidence, it was proposed that inactive 50S and 30S ribosomal subunits are nicked by RNase E and that rRNA fragments are then degraded by PNPase and RNase R (Zundel et al. 2009; Basturea et al. 2011; Sulthana et al. 2016). There is precedent in the literature for rifampicin-induced rRNA degradation. Zundel et al. (2009) have shown that addition of rifampicin to cells starved for glucose increased rRNA degradation. A recent report has shown that starvation for carbon, amino acids or phosphate as well as rifampicin treatment results in the degradation of rRNA (Fessler et al. 2020). Our work shows that rRNA degradation is a direct consequence of the inhibition of transcription resulting in the degradation of mRNA and the accumulation of inactive ribosomal subunits.

We have tested a battery of mutant strains lacking or deficient in enzymes involved in RNA degradation as well as a strain in which 10 toxin-antitoxin (TA) elements encoding mRNA interferases were deleted. None of the single gene deletions or the deletion of the TA elements affected rifampicin-dependent rRNA degradation. Our results with temperature-sensitive mutant strains affecting the activities of RNase E, PNPase and RNase R showed that these ribonucleases are involved in rRNA degradation. Rifampicin-induced rRNA degradation therefore follows the same pathway described previously for rRNA degradation during glucose starvation (Zundel et al. 2009; Basturea et al. 2011; Sulthana et al. 2016).

Nevertheless, our kinetic analysis in Fig. 7E and its quantification raises the issue of whether the effect of the RNase E temperature-sensitive mutants is direct or indirect. Since the inactivation of RNase E slows mRNA degradation, the lag in rRNA degradation could be due to a lag in the accumulation of inactive ribosomal subunits. Although not proof, we prefer the scenario in which RNase E acts directly in rRNA degradation since there is no evidence in Fig. 7 that other endo-ribonucleases such as RNase G, RNase III, or TA encoded mRNA interferases are involved in rifampicin-induced rRNA degradation.

The work reported here has implications for protocols that use inhibition of transcription to measure rates of mRNA degradation. 5S rRNA, which is stable under the conditions reported here, has often been used as an internal standard to normalize transcript levels in Northern blotting experiments. Whether there are other stable RNAs that can be reliably used as an internal standard is a question that should be explored. This is especially the case for genome-wide measurements of mRNA stability involving the synthesis of cDNA and high-density sequencing. The significant decrease in total RNA due to rifampicin-induced rRNA degradation reported here strongly suggests that normalizations assuming total RNA levels are constant will introduce a bias that results in an underestimation of mRNA degradation rates.

An equally important issue is that RNase E, PNPase and RNase R are major enzymes in rifampicin-induced rRNA degradation as well as mRNA degradation (Deutscher 2006; Carpousis et al. 2009; Mackie 2013; Hui et al. 2014). Upon addition of rifampicin, the degradation of 23S and 16S rRNA, which are highly abundant, very likely competes with the degradation of mRNA and thus results in a slowdown in mRNA degradation. As we know of no way to measure degradation rates without inhibiting mRNA synthesis, we believe that it will be necessary to model the effect of rRNA degradation on mRNA degradation rates and apply a correction that estimates degradation rates under normal growth conditions.

## Materials and Methods

### Culture medium, strains and cfu measurements

LB, MOPS media and agar plates (LA) were prepared as described (Miller 1972; Neidhardt et al. 1974). Glucose, glycerol or succinate (0.5%) and casamino acids (0.2%) were added to MOPS medium as indicated. Strains used in this work are listed in Tbl. 1. Biological replicates were performed by inoculating an overnight culture with a single colony from a freshly streaked LA plate and then diluting the overnight culture 1/100 and growing to OD_600_ = 0.5. RNA synthesis was inhibited with rifampin (150 µg/ml). Protein synthesis was inhibited with chloramphenicol (125 µg/ml) or kasugamycin (1 mg/ml). To measure colony forming units after treatment with rifampicin, **s**amples were serial diluted at room temperature in BU (per liter: 7 g NaH_2_PO_4_.2H_2_O, 4 g NaCl, 3 g KH_2_PO_4_) and plated on LA.

**Table 1.**
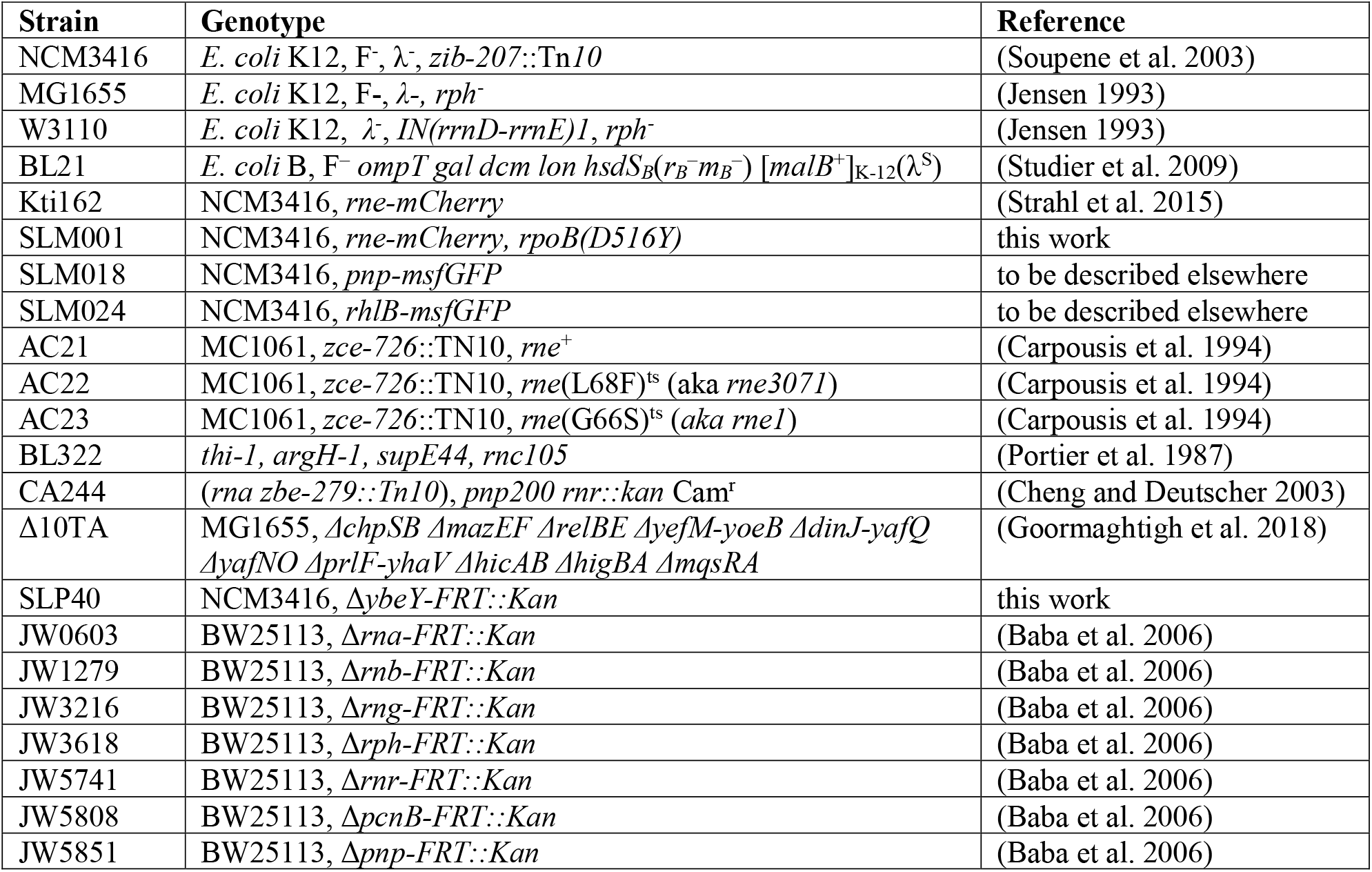
Strains.

### Measurement of cell dimensions

Overnight cultures were diluted in LB to an OD_600_ = 0.01 and grown at 37°C with vigorous shaking to an OD_600_ = 0.5. Cells were imaged before and 30 minutes after adding rifampicin (150 µg/ml). Cells (1-2 µl) were mounted on glass microscope slides with 1.2% (w/v) agarose pads. Acquisitions were made with a Nikon Eclipse TI-E/B wide field epifluorescence microscope using a phase contrast objective (CFI Plan Fluor DLL 100X oil NA1.3). Images were captured immediately at room temperature and edited using Nis-Elements AR software (Nikon). Images were analyzed using ImageJ v.1.38 software (National Institutes of Health) (Collins 2007; Schneider et al. 2012). Line scans were drawn over the long and short axis of cells to produce phase-contrast intensity profiles. Sharp intensity changes occurred where the lines crossed the boundary between the interior and exterior of the cell. Cell length and width were calculated from the distance between the points where the intensity change was half maximum (FWMH method, Full Width at Half Maximum). Statistical significance of the differences between untreated and rifampicin treated cells was calculated using the non-parametric Mann-Whitney test (GraphPad Prism version 8.0).

### Construction of isogenic rifampicin resistant strain

We obtained rifampicin resistant mutant strains by plating Kti162 on LA containing 200 µg/ml rifampicin. We selected a mutant strain for further characterization based on colony size and morphology comparable to the parent strain on LA plates at temperatures ranging from 20 to 43°C. DNA sequencing of the *rpoB* gene encoding the β-subunit of RNA polymerase showed that a D516Y amino acid substitution is responsible for rifampicin resistance. The aspartate residue at position 516 makes a direct contact with rifampicin (Campbell et al. 2001; Alifano et al. 2015). The mutation was genetically ‘purified’ by lambda RED recombineering using a 900 bp DNA fragment from the *rpoB* coding region and then phage P1 transduction to construct the SLM001 strain. Since the phenotypic expression of rifampicin resistance requires growth to replace wild type ribosomes with rifampicin resistant ribosomes, the P1 transduction protocol was modified by incubating phage transduced cells overnight before plating on rifampicin. In LB at 37°C, Kti162 and SLM001 have equivalent doubling times (21 min).

### Preparation of RNA, agarose gel electrophoresis and Northern blotting

Samples (5 ml) were taken from 75 ml cultures and quenched with 1 ml of 5% phenol (in ethanol) at room temperature. 0 minute time points were taken immediately after addition of rifampicin (150 µg/ml). After centrifugation (3000 g, 20 min, 4°C), supernatants were decanted, residual liquid was removed with a Kimwipe, and cell pellets were stored at -20°C. Total RNA was extracted using a Direct-zol™ RNA MiniPrep Plus kit (Zymo Research) as described (Hadjeras et al. 2019) except the DNase step was omitted. Briefly, cells were lysed under strongly denaturing conditions in TRIzol™. RNA was column purified and eluted in 50 µl of water. RNA amounts were quantitated by UV absorption using a Nano Drop™ spectrophotometer.

RNA was fractionated on mini format 2.4% agarose gels in 1x TBE at 100 V for 40 minutes. Samples were either mixed directly with a 2x formamide loading buffer or dried in a SpeedVac and suspended in 1x formamide loading buffer. Before loading, samples were heated in a ThermoMixer (50°C, 720 rpm, 5 min). Gels were stained with SYBR™ Safe and imaged with a GelDoc™ system. Northern blotting was performed as described (Hadjeras et al. 2019).

rRNA and tRNA species were quantified using Image Lab™ software. In Figs. 2, 4, 6 and 7, nominal amounts of RNA were determined using known quantities of the 750 and 1500 bp double-stranded DNA size markers as standards (1kb ladder, ThermoScientific). In these figures, the quantification is presented as a log_2_ plot to better visualize RNA levels that vary as much as 50-fold in some measurements.

## Acknowledgments

This work was supported by a grant from the Agence Nationale de la Recherche (IB-mRND, ANR-16-CE12-0014-02). We thank Jerome Rech for technical assistance on the Nikon microscope and Manual Campos, Laurence Girbal and Muriel Cocaign-Bousquet for helpful discussions and critical comments.

## Notes

### Competing Interest Statement

The authors have declared no competing interest.

